# Validation of Reference Genes for RT-qPCR in Marine Bivalve Ecotoxicology: Systematic Review and Case Study

**DOI:** 10.1101/074542

**Authors:** Moritz Volland, Julián Blasco, Miriam Hampel

## Abstract

Reverse transcription real-time quantitative PCR (RT-qPCR) is the predominant method of choice for the quantification of mRNA transcripts of a selected gene of interest. Here reference genes are commonly used to normalize non-biological variation in mRNA levels and their appropriate selection is therefore essential for the accurate interpretation the collected data. In recent years the use of multiple validated references genes has been shown to substantially increase the robustness of the normalization. It is therefore considered good practice to experimentally validate putative reference genes under specific experimental conditions, determine the optimal number of reference genes to be employed, and report the method or methods used.

Under this premise, we assessed the current state of reference gene base normalization in RT-qPCR bivalve ecotoxicology studies (post 2011), employing a systematic quantitative literature review. A total of 52 papers published met our criteria and were analysed for the gene or genes used, whether they employed multiple reference genes, as well as the validation method employed. In addition we performed a case study using primary hemocytes from the marine bivalve *Ruditapes philippinarum* after *in vitro* copper exposure. Herein we further critically discuss methods for reference gene validation, including the established algorithms geNorm, NormFinder and BestKeeper, as well as the popular online tool RefFinder.

We identified that RT-qPCR normalization in bivalve ecotoxicology studies is largely performed using single reference genes, while less than 40% of the studies attempted to experimentally validate the expression stability of the reference genes used. 18s rRNA and β-Actin were the most popular genes, yet their un-validated use did introduce artefactual variance that altered the interpretation of the resulting data, while the use of appropriately validated reference genes did substantially improve normalization. Our findings further suggest that combining the results from multiple individual algorithms and calculating the overall best-ranked gene, as e.g. computed by the RefFinder tool, does not by default lead to the identification of the most suitable reference gene or combination of reference genes.

## Introduction

Gene expression analysis is a powerful tool to investigate the underlying mechanisms of toxicant exposure in non-target organisms (as reviewed by Snape et al., 2004; Snell et al. 2003). Over the past three decades bivalve molluscs, such as the marine clam *Ruditapes philippinarum* (*R. philippinarum*), have been used as sentinel organisms to assess the potential effects of xenobiotics (in-depth reviewed by Cajaraville et al., 2000; Galloway and Depledge, 2001) and recently gene expression analysis has been employed to study potential mechanisms of pathogen and xenobiotic toxicity in these organisms (Brulle et al., 2012; Chen et al., 2015; Jeffroy et al., 2013; Liu et al., 2014; Liu et al., 2013; Miao et al., 2014; Milan et al., 2013; Moreira et al., 2012; Volland et al., 2015).

Reverse transcription real-time quantitative PCR (RT-qPCR) has been established as a method of choice for the quantification of mRNA transcripts of a selected gene of interest (GOI) in biological samples (as discussed by Bustin et al., 2009; Pfaffl et al., 2004; Wong and Medrano, 2005). Here reference genes (RGs) – colloquially referred to as housekeeping genes - are commonly used to normalize potential non-biological variance in mRNA levels introduced by sample handling and processing errors. In theory a RG is expressed in a wide variety of cells and tissues, and exhibits no variation in expression in response to experimental conditions. In reality, however, such a RG is unlikely to exist (Thellin et al., 1999). As variation in the RG expression can obscure real changes or produce artefactual ones, appropriate RG selection is essential for the accurate interpretation of RT-qPCR results (Bustin et al., 2009; Pfaffl et al., 2004; Wong and Medrano, 2005). The advantage of combining multiple validated reference genes into a so-called Normalization Factor (NF) has been established (Vandesompele et al., 2002) and the potential bias introduced by single gene normalization demonstrated (Tricarico et al., 2002). Consequently it is considered good laboratory practice to experimentally determine the optimal number of reference genes and to report the method or methods used, as defined by the *Minimum Information for Publication of Quantitative Real-Time PCR Experiments* (MIQE) guidelines (Bustin et al., 2009).

Unfortunately qPCR normalization in ecotoxicology studies using non-model organisms, such as bivalves, has widely been performed using single RGs, frequently with not validated genes or with genes previously validated under different experimental conditions; consequently making the interpretation of presented results difficult.

The aim of the present study was to review the current state of reference gene based normalization in RT-qPCR bivalve ecotoxicology studies, employing a systematic quantitative literature review. We further discuss reference gene validation methods, based on a case study assessing the expression stability of six common putative bivalve RGs in *R. philippinarum* hemocytes after various distinct *in vitro* copper challenges, including copper oxide (CuO) nano- and bulk particles, as well as ionic Cu (as CuSO_4_). Three well-established and widely used algorithms – geNorm (Vandesompele et al., 2002), BestKeeper (Pfaffl et al., 2004) and NormFinder (Andersen et al., 2004) – were used and our findings are contrasted to the common procedure of assessing RG expression stability by considering variability in the quantification cycles (Cqs). In addition the results from the individual algorithms are compared to RefFinder (Xie et al., 2012) a web-based tool that intents to combine the previous mentioned algorithms, as well as includes a forth method termed ∆Cq-method. We further outline a procedure to calculate global control group variability post normalization, in order to discuss the reference genes and their combinations proposed by the various methods presented in this study.

## Materials and method

### 1. Systematic quantitative literature review

A literature search was conducted on May the 9^th^ 2016 in Science Direct and ISI Web of Science using the following key words: “bivalve” and “gene expression”. We purposely selected peer-reviewed papers published after 2011 to reflect the current state of conduct and allow for the possible adoption of the MIQE guidelines (Bustin et al., 2009), while equally allowing for the consideration of a sufficient quantity of studies for a robust analysis. The returned selection (>1000 papers), was manually revised and studies meeting the following additional criteria were included in downstream analyses: i.) they were conducted on bivalves; ii.) they utilized RT-qPCR as a measure to study either biological marker genes (following stress exposure) or to validate such genes observed in high throughput mRNA studies; iii.) they did not focus on molecular characterization or cloning; and iv.) they employed one or a multitude of genes for the normalization of measured GOIs. Subsequently journal metrics were recorded and studies published in journals with 5-Year Impact Factors ≤2, according to the 2015 Journal Citation Reports*®* (Thomas Reuters, 2016), were excluded.

A total of 52 studies meeting our criteria were assessed as follows: i.) gene or genes employed for normalization; ii.) use of a single reference gene or the combination of multiple genes into a Normalization Factor; iii.) assessment of putative reference gene expression stability prior to normalization; here studies were divided into either using an experimental assessment process, providing references to prior work performed, or providing neither; iv.) experimental validation strategies employed; and v.) disclosure of results from the experimental expression stability assessment.

Studies including statements regarding the stability and/or variability of only one putative reference gene without providing some indication as to how this conclusion was reached or providing data on the variability were not counted as utilizing an experimental assessment process. Studies partially including expression stability assessment results (e.g. providing geNorm M values, but no V values) were counted as disclosing results. All quantitative reviewed research and data extracted have been included in the Supplementary Materials (Table SM1).

### 2. Treatments

CuO NPs with nominal similar size distributions of 25 - 55 and ~40 nm (from here on called CuO_25-55 and CuO_40, respectively) and similar purity (>99.95% and >99% respectively) were purchased from US Research Nanomaterials (Houston, USA) (product numbers US3063 and US3070, respectively). Bulk copper oxide particles (bCuO) and ionic copper (as CuSO_4_) of equal purity (99.99%) were purchased from Sigma-Aldrich (Madrid, Spain) (product numbers 450812 and 451657, respectively). All copper forms were purchased as powders. Additional particle characterization has been included in the Supplementary Materials (Section 1).

### 3. Hemocyte collection, culture and *in vitro* exposure

Adult clams (*Ruditapes philippinarum*) (size: 3 - 4.5 cm, mean weight +/- SD: 13.6 +/-3.0 g) were supplied by a commercial clam fishery (Mariscos Ria de Vigo S.L., Vigo-Pontevedra, Spain). Clam hemocyte isolation and culture was based on the protocol by Gomez-Mendikute and Cajaraville (2003), as modified by Katsumiti et al. (2014). Hemolymph was withdrawn under aseptic conditions from the posterior and anterior adductor muscle. Per independent replicate the hemolymph of 18-22 animals was pooled and diluted to 8 × 10^5^ cells mL^−1^ (>95% viable according to trypan blue exclusion assay). Per exposure condition and replicate 200 µL cell suspensions were seeded into six wells of a 96-well microplate in culture medium. Subsequently microplates were centrifuged at 270 g for 10 min at 4° C in order to assist cell attachment and cells were maintained for 24 h prior to exposure. Freshly dispersed suspensions in cell culture medium were prepared (1 mg Cu mL^−1^), sonicated for 10 min, serial-diluted to appropriate concentrations (1 and 10 µg Cu mL^−1^), and immediately applied to the cells. Exposures were performed for 24 h and cells not exposed to particles served as negative controls.

### 4. Relative gene expression analysis

#### 4.1. Hemocyte collection, RNA extraction and cDNA synthesis

Hemocytes from 4 independent replicates per treatment (n=4) and 8 independent replicates for the control group (n=8) were collected by re-suspension via gentle pipetting of cells in wells. Following centrifugation at 500 g for 10 min at 4° C, cells were washed twice with ice-cold phosphate-buffered saline (PBS, pH 7.4), re-suspended in 100 µL PBS and added to 0.5 mL RNAlater^®^ (Sigma Aldrich). Prior to RNA extraction in Tri Reagent^®^ applying the Sigma^®^ standard protocol (Sigma Aldrich) cells were pelleted at 2500 g for 5 min at 4° C and washed with PBS (4° C). Extracted RNA was purified using the NucleoSpin^®^ RNA XS kit (Macherey-Nagel, Düren, Germany), according to the manufacturer protocol. RNA quantity was measured in a Qubit^®^ 2.0 Fluorometer using a Qubit^®^ RNA Assay Kit (Thermo Fischer Scientific), and its quality was determined in a Bioanalyzer 2100 with the RNA 6000 Nano kit (Agilent Technologies, Santa Clara, USA) following the method described for invertebrate RNA by Winnebeck et al. (2010). Total RNA (~100 ng) was reverse-transcribed using the QuantiTect^®^ Reverse Transcription Kit (Qiagen, Venlo, Netherlands) following the standard protocol. For the relative quantification of gene expression using real-time PCR (qPCR) of selected genes (Table 1 and Supplementary Materials Table SM2), specific primer pairs were designed using Primer 3 version 4.0.0 software (Untergasser et al., 2012) (available at http://bioinfo.ut.ee), unless stated otherwise (Table 1 and Supplementary Materials Table SM2), and their oligonucleotide sequences were synthesized by Biomers^®^ (Ulm, Germany). qPCR reactions were performed on a Mastercycler^®^ epgradient S Realplex2 with Realplex software version 2.2 (Eppendorf, Hamburg, Germany). Assay linearity and amplification efficiency were established (0.991-0.998 and 0.86-1.04 respectively), as well as primer concentration (200 nM). Melting curve analysis was used to ensure primer specificity. Annealing temperature was set to 60 °C. Assay linearity and amplification efficiency of primer pairs have been included in the Supplementary Materials Table SM2. Samples were removed from downstream analysis i.) when Bioanalyzer 2100 profiles were not as previously described (Winnebeck et al., 2010); ii.) less than 100 ng total RNA could be isolated; or iii) the standard deviation between technical replicates was >0.15 quantification cycles (Cqs). Additional information related to RNA extraction, cDNA synthesis, and RT-qPCR conditions, as well as raw Cq values have been included in the Supplementary Materials (Section 2, Table SM3).

**Table 1.**
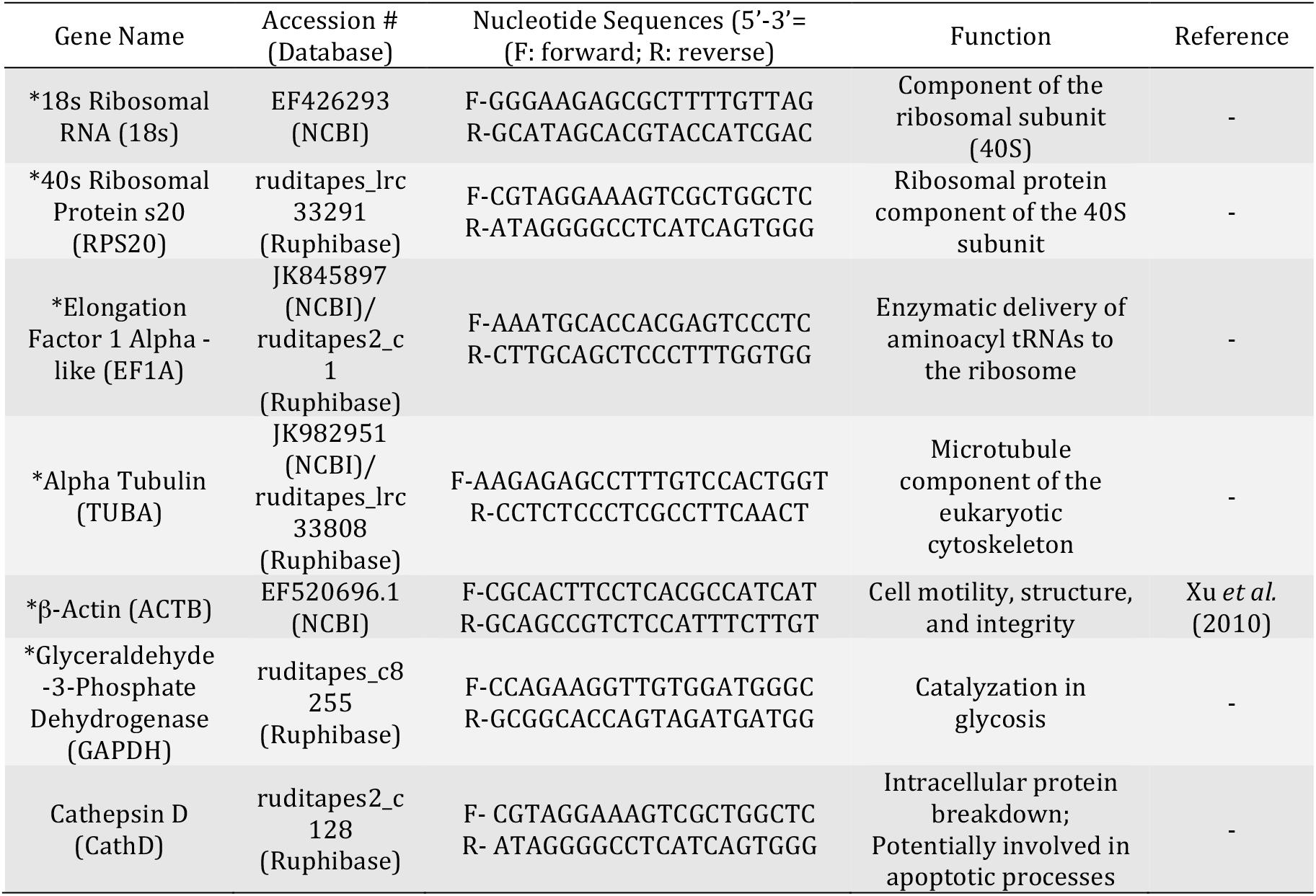
Description of the genes and primers used in this study. * Marked genes were used as putative reference genes.

#### 4.2. Relative expression stability analysis of candidate reference genes

A panel of 6 common putative RGs was included in the study: 18s ribosomal RNA (18s), Glyceraldehyde 3-Phosphate Dehydrogenase (GAPDH), β-Actin (ACTB), a-Tubulin (TUBA), Elongation Factor 1 a – like (EF1A), as well as 40S Ribosomal Protein S20 (RPS20). Using three common and distinct algorithms - geNorm (Vandesompele et al., 2002), NormFinder (Andersen et al., 2004) and BestKeeper (Pfaffl et al., 2004) - their relative expression stability was assessed under experimental conditions. In addition results were contrasted to the RefFinder program (Xie et al., 2012) a popular web-based tool for RG selection. The geNorm algorithm was used through the R NormqPCR packages (Version 1.17.0) (Perkins et al., 2012). The NormFinder algorithm was implemented as an R-script (NormFinder for R version 5, 2015-01-05; available at http://moma.dk/normfinder-software). We set NormFinder sub-groups to consider all treatment-concentration combinations and due to concerns over sub-group size (3-4 replicates per sub-group) influencing model accuracy, a contrast considering only treatment groups was tested as well (6-8 replicates per subgroup). The BestKeeper algorithm, used through the BestKeeper excel tool (Version 1), computes a summary statistic of the Cq values for each candidate gene, as well as a Pearson correlation coefficients with the BestKeeper index (geometric mean of all included genes), with a resulting coefficient close to 1 being desired. Genes were preliminarily ranked based on the calculated Cq standard deviations (SD) and unstable genes (SD>1) were excluded from the BestKeeper index, as suggested by the authors. RefFinder (http://fulxie.0fees.us/?type=reference) intents to integrate the previous listed algorithms and includes a consensus ranking, based on the geometric mean of the individual ranks of each implemented method. Additional details on the different algorithms have been included in the Supplementary Materials (Section 3).

#### 4.3. Data analysis and relative quantification of gene expression

Statistical analysis was performed using R version 3.2.4 (R Core Team, 2016) including the following packages; raw Cq values were read with the ReadqPCR package (Version 1.17.0) (Perkins et al., 2012) and ∆Cqs were calculated, as described below (formula [1]), using the NormqPCR package, both available through the Bioconductor project (Version 3.2) (Huber et al., 2015), and data were graphed using the ggplot2 package (Wickham, 2009). For ease of visual comparison data are presented as relative expression profiles - log2 fold changes (∆∆Cqs) - after normalization against a calibrator (control samples), calculated as shown below in formulas [1] - [3]:

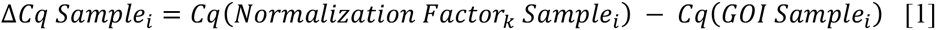

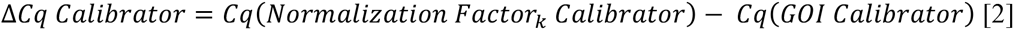

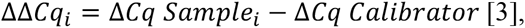

where the Normalization Factor *k* (*NF*_*k*_) was a single putative RG or the arithmetic mean from two or more putative RGs (formula [4]):

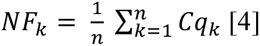

#### 4.4. Relative normalization control error

Under the assumptions that there is no systematic variation within the control group and that variation in a selected Normalization Factor (NF) (formula 9) will introduce artefactual variation in the relative quantities of normalized gene expression, we analysed global expression variability within the control group post normalization for 9 GOIs included in the study (Supplementary Materials Table SM2) as a proxy for resulting normalization error. This overall variability, here called Normalization Control Error (*NCE*), can be calculated independent of relative quantification strategies as shown below in formulas [5] – [7]:

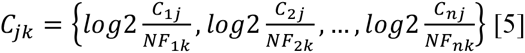

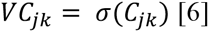

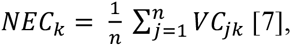

where *C* is the relative expression of gene of interest *j* (formula [8]) and NF the geometric mean of the relative expression of putative RGs included in the Normalization Factor *k* (formula [9]):

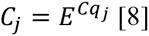

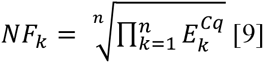

Note that for relative quantification strategies where primer efficiencies are assumed to be 100% (E=2) *C*_*jk*_ can be calculated as shown in formula [10]. Here the Normalization Factor is calculated as previously shown in formula [4]:

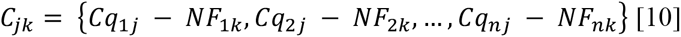

## Results

### 1. Current use of reference genes in RT-qPCR bivalve studies

A total of 52 peer-reviewed papers, with an average 2015 Journal Citation Reports^®^ 5-Year Impact Factor of 3.57 (ranging from 2.21 to 5.60), were assessed. As can be seen in Figure 1A, the majority of studies reviewed used single reference gene (RG) normalization, with only 13.7% combining multiple genes into a Normalization Factor. The most common reference genes were 18s ribosomal RNA (rRNA) and β-Actin (ACTB), with 22 and 15 citations respectively, followed by Elongation Factor-α (EFA) > 28s rRNA > Glyceraldehyde-3-Phosphate Dehydrogenase (GAPDH) ≥ Ribosomal Protein L27 (RPL27) ≥ Tubulin-α (TUBA) ≥ Cytochrome C Oxidase 1 (CO1) > Peroxiredoxin (PRXD) ≥ Ribosomal Protein S3 (RPS3) ≥ Sulfotransferase (SULT) ≥ Tubulin-β (TUBB) (Figure 1B). Less than 40% of studies attempted to experimentally validate the RG or RGs used under the experimental conditions (Figure 1C), while of those less than 25% presented the results of said assessments (data not shown). An additional 15.4% provided reference to previous research performed that employed the utilized gene or genes. The most popular methods to assess expression stability were geNorm and Cq sample variability, here termed Cq Variance, followed by BestKeeper and Normfinder, while in some cases the selected method was not clearly identifiable (Figure 1D).

**Figure 1.**
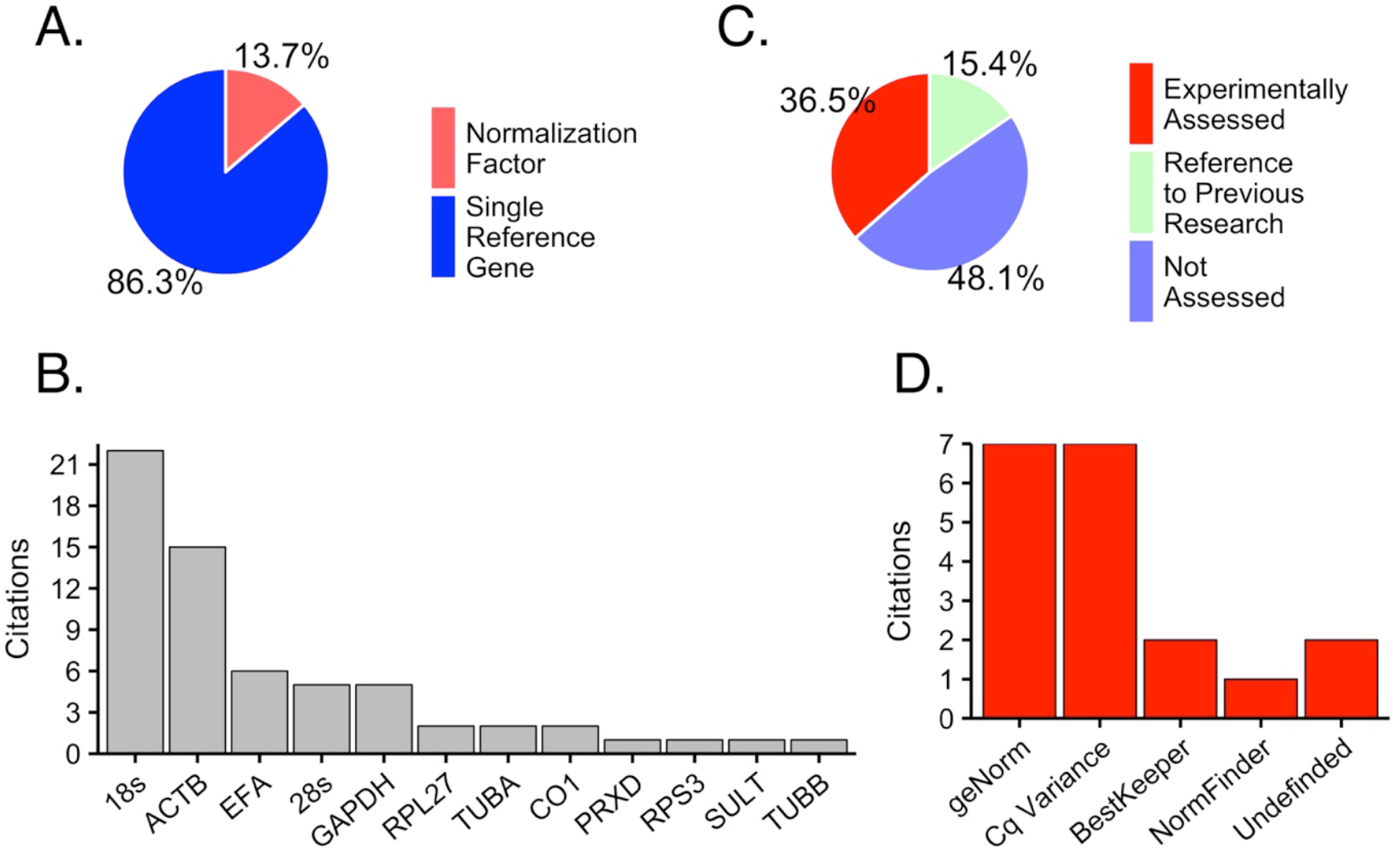
Summary data on the use of reference genes in RT-qPCR biomarker analyses in 52 bivalve studies published between 01^st^ January 2012 and 9^th^ of May 2016. (A.) Percentage of studies using a single reference gene or the combination of multiple genes (Normalization Factors) for RT-qPCR normalization. (B.) Genes cited as reference genes. (C.) Percentage of studies performing experimental assessment of reference genes (‘Experimentally Assessed’), provide reference to previous research employing the utilized reference genes (‘Reference to Previous Research’) or providing neither (‘Not Assessed’). (D.) Algorithms cited as employed for the experimental validation of reference gene.

### 2. Health status hemocytes

Following the various copper treatments cell viability was, in parts, compromised by specific treatments, as quantified compared to the control group (100%). Following the exposure to 1 and 10 ppm (in Cu) respectively, CuO_25-55 reduced cell viability by 4.1 and 28.8%, bCuO by 5.2 and 9.3%, CuSO_4_ by 2.2 and 6.4%, and CuO_40 by 2.6 and 4.8%. Control samples had a cell viability of >95% in all performed assays, as established by the trypan blue assay.

### 3. RNA quality

As previously reported protostome 28s rRNA frequently fragments into two similarly sized fragments during routine RNA extraction procedures. As these fractions tend to migrate comparable to 18s during electrophoresis, the Bioanalyzer (Agilent) RNA integrity numbers cannot be used to assess RNA quality (Winnebeck et al., 2010). Instead electrophoresis profiles were used to visually assess degradation as previously described by Winnebeck and colleagues (2010).

### 4. Gene expression and expression variability

We calculated the mean Cq value and standard deviation for each gene (n=34). EF1A (Cq=18.6) was the most expressed RG, followed by TUBA (Cq=19.5), 18s (Cq=21.5), RPS20 (Cq=21.7), GAPDH (Cq=23.1) and ACTB (Cq=24.8). As shown in Figure 2, RPS20 exhibited the smallest range in Cq values, as well as the smallest standard deviation (0.6), followed by TUBA (0.7), EF1A (0.8), ACTB (1.0), GAPDH (1.1), and 18s (2.2).

**Figure 2.**
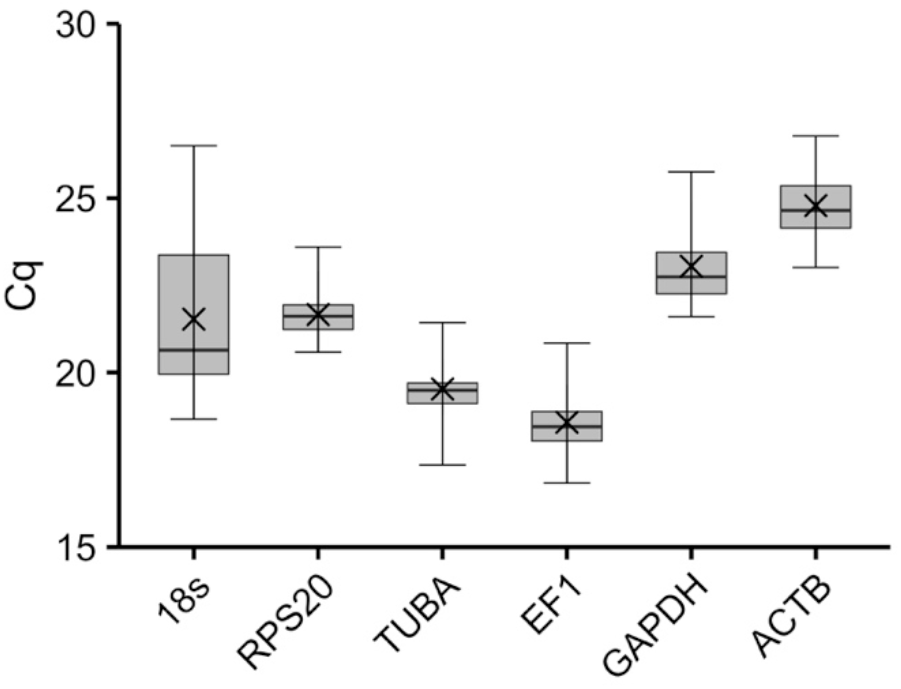
RT-qPCR cycle threshold (Cq) summary for 6 putative reference genes in 34 *R. philippinarum* haemocyte RNA samples. The median Cqs are shown as horizontal lines, mean Cqs are shown as (X), 25 to 75 percentiles are included as the shaded boxes and the full range of Cq values are given by the whiskers.

### 5. Quantitative analysis of candidate reference genes based on BestKeeper, geNorm, NormFinder and RefFinder

The different algorithms identified, in parts, different reference genes or pairs of genes for optimal normalization, while uniformly 18s was identified to be the least stable expressed gene. geNorm identified RPS20 and TUBA to be the most stable RGs, followed by EF1A, GAPDH, ACTB and ultimately 18s (Figure 3A, Table 2). The calculated pairwise variation value for the inclusion of EF1A as an additional RG was below the suggested cut-off value (0.15) (Figure 3B), suggesting that its inclusion would not significantly alter the normalization. Except 18s, all candidate reference genes could be considered stable for a homogeneous sample group with geNorm stability values (M) below 0.5 (Hellemans et al., 2007). NormFinder identified GAPDH to be the most stable RG tested, followed by EF1A, ACTB, TUBA, RPS20 and 18s. The assessment of the pairwise stability of two RGs identified GAPDH and EF1A to be the best pair, while the calculated stability was better than for single gene normalization (Figure 4, Table 2). In agreement with previous results, BestKeeper calculated standard deviations suggested the removal of 18s from the BestKeeper Index (BK). Subsequent Pearson correlations between candidate reference genes and the BK was highest for EF1A, followed by GAPDH, TUBA, RPS20 and ACTB (Table 2). As BestKeeper does not provide a measurement for the optimal number of RGs, we included the three genes with the highest correlation coefficient, as suggested by the authors as the minimal number of RGs (Pfaffl et al., 2004). RefFinder ranked EF1A as the best choice RG, followed by RPS20, TUBA, GAPDH, ACTB and 18s (Supplementary Materials Table SM3). Due to program specific limitations RefFinder calculated in parts different rankings for NormFinder and BestKeeper algorithms, as for BestKeeper only SDs were calculated and for NormFinder no sub-groups could be specified (Table 2 and Table SM3).

**Figure 3.**
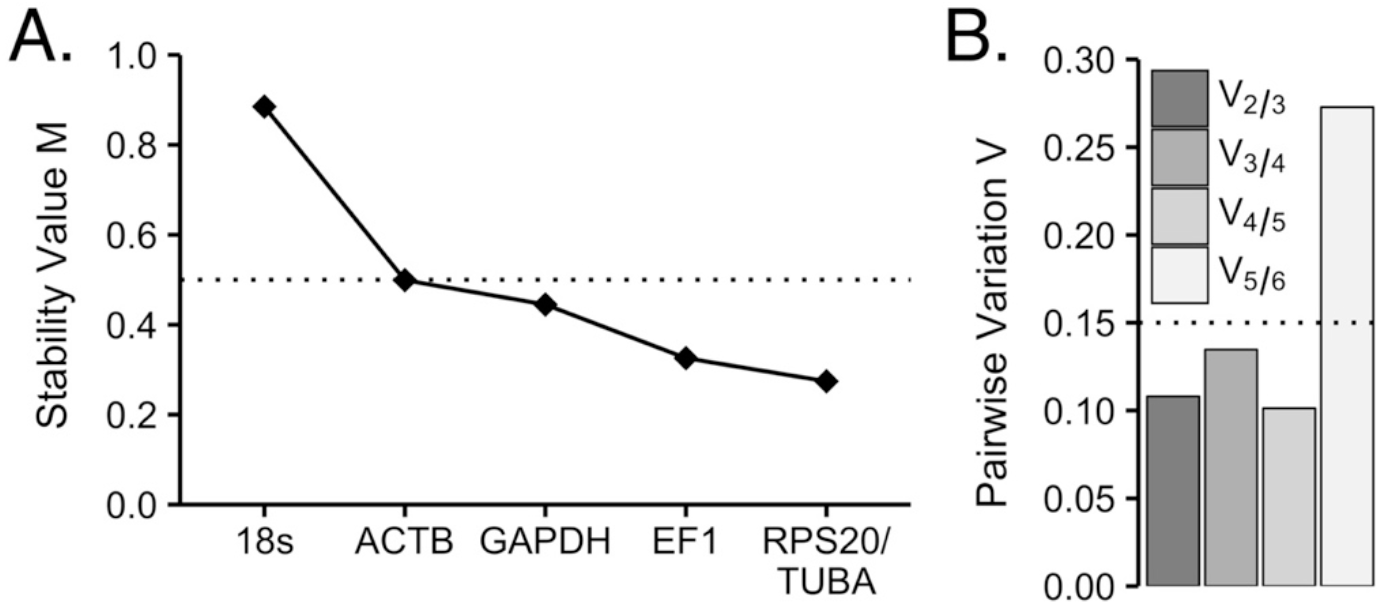
geNorm reference gene expression stability assessment. (A.) Expression stability values (M) of 6 putative reference genes after stepwise exclusion of the least stable expressed genes. Lower values denote higher stability. The dotted horizontal line represents the proposed cut-off value for unstable genes (>0.5). (B.) Pairwise variations analysis showing optimal number of reference genes to be included in the Normalization Factor. Pairwise variation values (V) below 0.15 (dotted horizontal line) indicate that the inclusion of an additional reference gene is not expected to substantially influence normalization.

**Figure 4.**
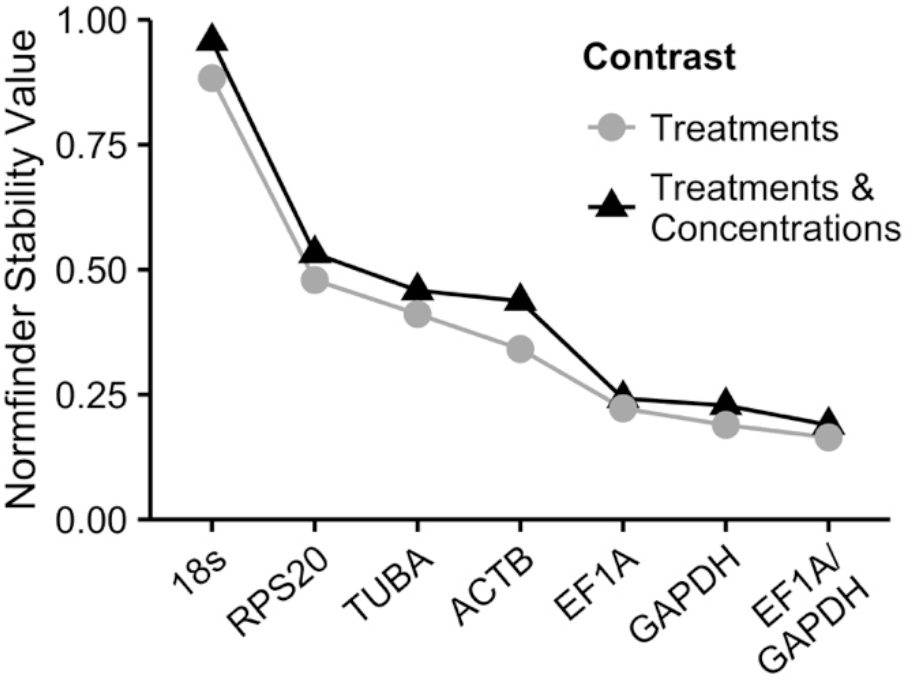
NormFinder gene expression stability for 6 putative reference genes and the pairwise stability of the combination of the two most suitable genes (EF1A and GAPDH). Lower values denote higher stability. Differing point shapes and corresponding line colours denote different contrasts (sub-groups) tested. The black line and triangles represent data from the analysis with a contrast consisting of all treatment and concentration combinations (nine sub-groups) and the grey line and dots represent data from the analysis with a contrast consisting of all distinct treatments (five sub-groups).

**Table 2.** Expression stability values of the candidate reference genes calculated by the BestKeeper, NormFinder and geNorm algorithms. Best ranked value(s) for each algorithm is (are) given in bold. Best NormFinder pairwise stability values are given in parentheses. ^a^: Values closer to 1 denote higher stability, ^b^: Values closer to 0 denote higher stability, ^c^: Value above suggested cut-off value for stable expressed genes for the respective method.

| Gene | BestKeeper |  | NormFinder | geNorm |
| --- | --- | --- | --- | --- |
|  | [R]<sup>a</sup> | SD<sup>b</sup> | Stability Value<sup>b</sup> | M Value<sup>b</sup> |
| 18s | - | 1.897<sup>c</sup> | 0.957 | 0.885<sup>c</sup> |
| RPS20 | 0.942 | **0.461** | 0.532 | **0.274** |
| TUBA | 0.954 | 0.503 | 0.458 | **0.274** |
| ACTB | 0.919 | 0.823 | 0.437 | 0.499 |
| EF1A | **0.973** | 0.640 | 0.242 (**0.189**) | 0.326 |
| GAPDH | 0.962 | 0.867 | **0.228 (0.189)** | 0.445 |

### 6. Relative normalization control variability

Under the assumption that sub-optimal normalization would produce artefactual variance in the control group, we ranked the success of various normalization strategies by their relative control group variability, here termed Normalization Control Error (NCE), as calculated for 9 genes of interest (n=54). As shown in Figure 5, in agreement with the employed algorithms single gene normalization by 18s and ACTB produced the greatest variability. In addition, single gene normalization with EF1A as identified by the RefFinder algorithm ranked worse than the NFs calculated by the individual algorithms. While the inclusion of the two best ranked RefFinder genes into a NF reduced variability, its rank was still lower than for the NFs calculated by the individual algorithms. Further, single gene normalization based on the gene with the overall smallest Cq variance (RPS20), ranked worse than NFs identified by geNorm, BestKeeper and NormFinder.

**Figure 5.**
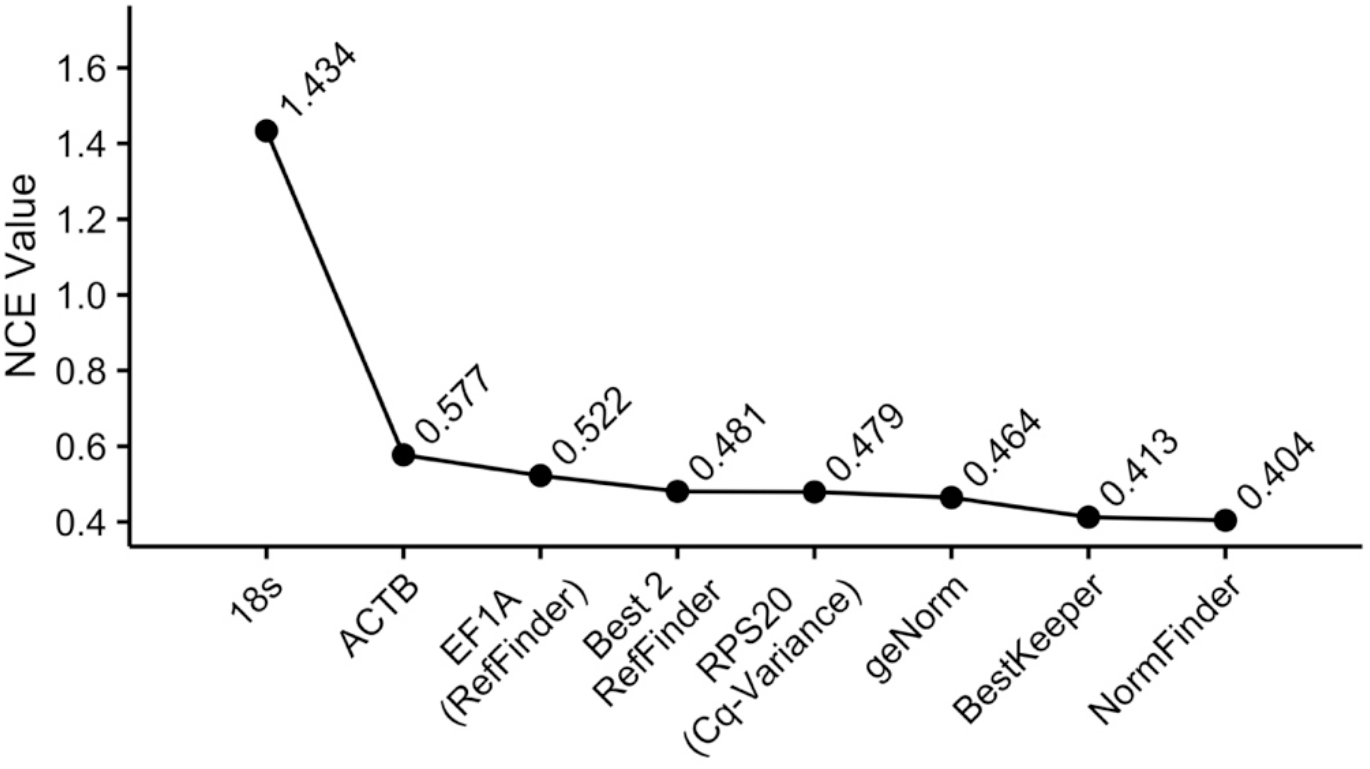
Normalization control error (NCE value), calculated as control group variance of nine target genes post normalization. Lower values denote higher stability. Respective normalization was performed with a single gene for 18s, ACTB, EF1A and RPS20 or with a Normalization Factor consisting of two or more genes as suggested by the various algorithms for geNorm (RPS20 and TUBA), BestKeeper (EF1A, GAPDH and TUBA) and NormFinder (EF1A and GAPDH), as well as for the best two ranked genes by the RefFinder tool (Best 2 RefFinder: EF1A and TUBA).

### 7. Relative expression profiles

As shown in Figure 6, expression profiles vary in parts substantially between normalization strategies. In agreement with the various algorithms employed the relative expression profile of 18s normalized Cathepsin D shows a very different profile than all other expression profiles.

**Figure 6.**
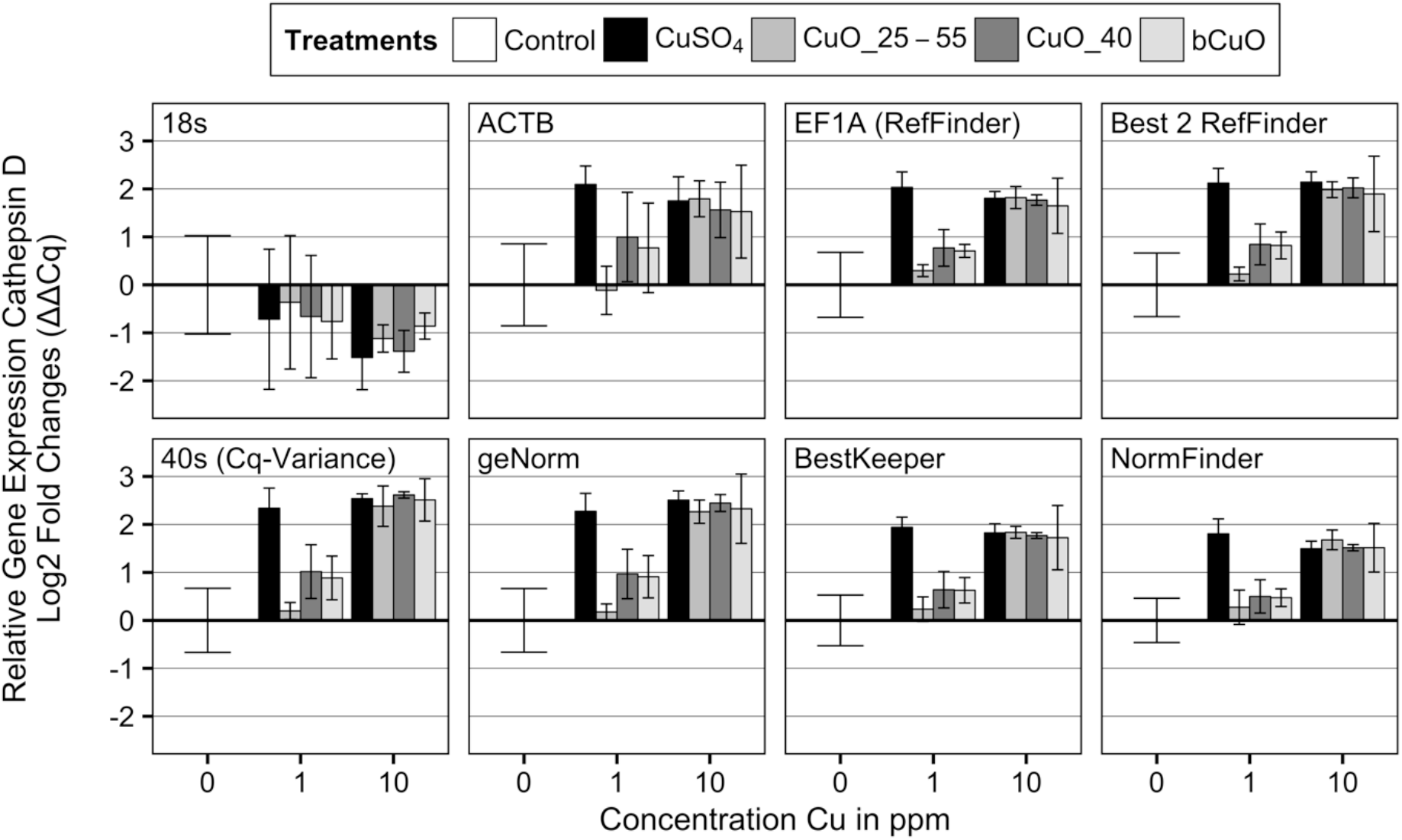
Relative Cathepsin D expression profiles after 24 h exposure to 0,1 and 10 ppm Cu of various copper treatments. Respective normalization was performed with a single gene for 18s, ACTB, EF1A and RPS20 or with a Normalization Factor consisting of two or more genes as suggested by the various algorithms for geNorm (RPS20 and TUBA), BestKeeper (EF1A, GAPDH and TUBA) and NormFinder (EF1A and GAPDH), as well as for the best two ranked genes by the RefFinder tool (Best 2 RefFinder : EF1A and TUBA).

## Discussion

Despite the existence of several established algorithms for the assessment of the expression stability of putative reference genes, RT-qPCR normalization in bivalves is commonly performed with single un-validated genes (Figure 1A). An approach that has been demonstrated to be prone to increased error (Vandesompele et al., 2002) and bias (Tricarico et al., 2002).

The two most common RT-qPCR RGs for the here reviewed studies were 18s rRNA and β-Actin (Figure 1B). Our results indicate, however, that both genes are sub-optimal RGs under the tested experimental conditions. The relative expression profiles for Cathepsin D (Figure 6) demonstrate that in particular normalization with 18s appears highly problematic for gene expression data derived from *R. philippinarum* hemocytes post copper exposure, largely altering the interpretation of the data. In agreement all tested algorithms ranked 18s the least stable among the tested candidate reference genes, being above the suggested acceptable geNorm stability value and BestKeeper standard deviation (M > 0.5 and SD > 1, respectively). β-Actin was ranked the second least stable candidate gene among the tested by the BestKeeper Pearson correlation analysis and geNorm algorithm (Table 2, Figure 3), as well as the here presented NCE (Figure 5). Our findings are in agreement with previous efforts in a number of bivalve species: despite their widespread use 18s (Araya et al., 2008; Du et al., 2013; Liu et al., 2013; Moreira et al., 2015; Wan et al., 2011) and β-Actin (Feng et al., 2013; Liu et al., 2013; Mateo et al., 2010; Moreira et al., 2015; Morga et al., 2010) have been frequently reported to be among the least stable assessed candidate reference genes.

Our results further indicate that EF1A, GAPDH, TUBA and RPS20 can, under the tested conditions, be used for accurate normalization of GOIs, in the various combinations suggested by the individual algorithms. While these genes have been frequently reported as the most stable reference genes in bivalve studies, both by themselves and in various combinations, no consensus can be reported (Araya et al., 2008; Du et al., 2013; Feng et al., 2013; Liu et al., 2013; Mateo et al., 2010; Moreira et al., 2015; Morga et al., 2010; Wan et al., 2011). Considering the, in parts, distinct underlying mechanisms and assumptions of the different algorithms used it is not surprising that for four relative constitutively expressed candidate genes algorithm-specific combinations were calculated.

Despite its popularity (Figure 1D), our data suggests that the exclusive assessment of Cq variability can be an insufficient approach for the selection of one or multiple RGs (Figure 5). To overcome the difficulty in choosing the optimal RG or combination of genes, a consensus approach has recently been employed in bivalve studies (Feng et al., 2013; Moreira et al., 2015). RefFinder is a popular tool for candidate reference gene identification (>10 citations) employing the consensus approach, as recently reviewed by De Spiegelaere et al. (2015). Here the results of the individual algorithms are ranked and the candidate reference gene with the overall best rank is proposed as the optimal reference gene. This approach however has several limitations: i.) as previously discussed single gene normalization is generally considered to be a sub-optimal approach (Bustin et al., 2009; Tricarico et al., 2002; Vandesompele et al., 2002; Wong and Medrano, 2005); ii.) combining two or more of the top ranked consensus genes to a NF without assessing their pairwise stability might transfer instead of eliminate the systematic error (Andersen et al., 2004); and iii.) no indication on the optimal number of genes to include can be inferred. In agreement with the above-discussed limitations Figure 5 shows that single gene normalization using the RefFinder identified consensus RG did not result in the most accurate normalization. While the combination of the two best-ranked genes to a NF (here termed *Best 2 RefFinder*) reduced the resulting variability somewhat, it was still greater than for the NFs suggested by the individual algorithms.

Considering that over 60% of the here reviewed studies did not experimentally validate the employed RGs (Figure 1C), it should be noted that candidate RG stability patterns in bivalves have been found to differ even within the same species between tissues and life stages (Feng et al., 2013) and after exposure to related bacteria strains (Mateo et al., 2010). It has further been demonstrated that varying pressures during seasonality, such as thermal variation, food availability and reproductive status (Banni et al., 2011; Hamano et al., 2005), as well as sex (Veldhoen et al., 2009) can affect gene expression patterns and up-to-date not enough evidence exists to exclude potential effects on the expression of putative reference genes. Apart from biological restrains when replicating experimental conditions, one should consider the statistical limitations of the employed algorithms. While RGs and/or NFs are generally ranked or cut-off values for the inclusion of additional genes into a NF are provided, statistical inference testing is not performed and consequently results should be considered as sample and not population specific. In agreement with previous researchers (Li et al., 2012; Wong and Medrano, 2005) our findings suggest that suitable reference genes should be established for each species, tissue or cell type, and specific experimental conditions to ensure their stability, instead of relying on previous research. Unfortunately this is still not the case in the majority of bivalve studies utilizing RT-qPCR to analyse molecular markers (Figure 1C). In addition we recommended to employ a minimum of two experimentally validation procedures, as our data shows that individual algorithms can produce varying suggestions.

## Conclusion

We conclude that 18s rRNA and β-Actin are, despite there widespread use, not appropriate reference genes for the normalization RT-qPCR data from *R. philippinarum* hemocytes post copper exposure, as well as frequently being identified among the least stable genes tested in bivalve studies. EF1A, GAPDH, TUBA and RPS20, in the various combinations suggested by geNorm, NormFinder and BestKeeper, were suitable for the normalization under the tested conditions and we therefore recommend their inclusion in the assessment of candidate reference genes for *R. philippinarum* hemocyte exposure studies. We further identified that the consensus approach, as for example employed by the RefFinder tool, does not by default lead to the selection of the most appropriate gene or genes for normalization. In addition, this approach does neither allow for the identification of the optimal number of reference genes, nor their optimal combination. Although several of the established algorithms and their software packages might at first appear inaccessible to researchers, a clear understanding of their strengths and weaknesses is necessary to accurately interpret the obtained results and subsequently decide on the most appropriate normalization strategy.

## Acknowledgments

This work was funded by the regional government of Andalusia (Junta de Andalucía) project PE2011-RNM-7812, as well as supported by the Erasmus Mundus Ph.D. fellowship in Marine and Coastal Management to M. V. (as coordinated by the University of Cadiz, Spain). The authors would further like to express their gratitude to Dr. Cajaraville and Dr. Katsumiti from the Department of Zoology and Animal Cell Biology at the University of the Basque Country UPV/EHU.

## References

Andersen, C.L., Jensen, J.L., Orntoft, T.F., 2004. Normalization of real-time quantitative reverse transcription-PCR data: a model-based variance estimation approach to identify genes suited for normalization, applied to bladder and colon cancer data sets. Cancer Research 64, 5245–5250. 10.1158/0008-5472.CAN-04-0496

Araya, M.T., Siah, A., Mateo, D., Markham, F., Mckenna, P., Johnson, G., Berthe, F.C., 2008. Selection and evaluation of housekeeping genes for haemocytes of soft-shell clams (Mya arenaria) challenged with Vibrio splendidus. Journal of Invertebrate Pathology 99, 326–331. 10.1016/j.jip.2008.08.002

Banni, M., Negri, A., Mignone, F., Boussetta, H., Viarengo, A., Dondero, F., 2011. Gene expression rhythms in the mussel Mytilus galloprovincialis (Lam.) across an annual cycle. PLoS One 6, e18904. 10.1371/journal.pone.0018904

Brulle, F., Jeffroy, F., Madec, S., Nicolas, J.L., Paillard, C., 2012. Transcriptomic analysis of Ruditapes philippinarum hemocytes reveals cytoskeleton disruption after in vitro Vibrio tapetis challenge. Developmental & Comparative Immunology 38, 368–376. 10.1016/j.dci.2012.03.003

Bustin, S.A., Benes, V., Garson, J.A., Hellemans, J., Huggett, J., Kubista, M., Mueller, R., Nolan, T., Pfaffl, M.W., Shipley, G.L., Vandesompele, J., Wittwer, C.T., 2009. The MIQE guidelines: minimum information for publication of quantitative real-time PCR experiments. Clinical Chemistry 55, 611–622. 10.1373/clinchem.2008.112797

Cajaraville, M.P., Bebianno, M.J., Blasco, J., Porte, C., Sarasquete, C., Viarengo, A., 2000. The use of biomarkers to assess the impact of pollution in coastal environments of the Iberian Peninsula: a practical approach. Science of the Total Environment 247, 295–311. 10.1016/s0048-9697(99)00499-4

Chen, H., Zha, J., Yuan, L., Wang, Z., 2015. Effects of fluoxetine on behavior, antioxidant enzyme systems, and multixenobiotic resistance in the Asian clam Corbicula fluminea. Chemosphere 119, 856–862. 10.1016/j.chemosphere.2014.08.062

De Spiegelaere, W., Dern-Wieloch, J., Weigel, R., Schumacher, V., Schorle, H., Nettersheim, D., Bergmann, M., Brehm, R., Kliesch, S., Vandekerckhove, L., Fink, C., 2015. Reference gene validation for RT-qPCR, a note on different available software packages. PLoS One 10, e0122515. 10.1371/journal.pone.0122515

Du, Y., Zhang, L., Xu, F., Huang, B., Zhang, G., Li, L., 2013. Validation of housekeeping genes as internal controls for studying gene expression during Pacific oyster (Crassostrea gigas) development by quantitative real-time PCR. Fish & Shellfish Immunology 34, 939–945. 10.1016/j.fsi.2012.12.007

Feng, L., Yu, Q., Li, X., Ning, X., Wang, J., Zou, J., Zhang, L., Wang, S., Hu, J., Hu, X., Bao, Z., 2013. Identification of reference genes for qRT-PCR analysis in Yesso scallop Patinopecten yessoensis. PLoS One 8, e75609. 10.1371/journal.pone.0075609

Galloway, T.S., Depledge, M.H., 2001. Immunotoxicity in invertebrates: measurement and ecotoxicological relevance. Ecotoxicology 10, 5–23. 10.1023/A:1008939520263

Gomez-Mendikute, A., Cajaraville, M.P., 2003. Comparative effects of cadmium, copper, paraquat and benzo[a]pyrene on the actin cytoskeleton and production of reactive oxygen species (ROS) in mussel haemocytes. Toxicology In Vitro 17, 539–546. 10.1016/S0887-2333(03)00093-6

Hamano, K., Awaji, M., Usuki, H., 2005. cDNA structure of an insulin-related peptide in the Pacific oyster and seasonal changes in the gene expression. Journal of Endocrinology 187, 55–67. 10.1677/joe.1.06284

Hellemans, J., Mortier, G., De Paepe, A., Speleman, F., Vandesompele, J., 2007. qBase relative quantification framework and software for management and automated analysis of real-time quantitative PCR data. Genome Biology 8, R19. 10.1186/gb-2007-8-2-r19

Huber, W., Carey, V.J., Gentleman, R., Anders, S., Carlson, M., Carvalho, B.S., Bravo, H.C., Davis, S., Gatto, L., Girke, T., Gottardo, R., Hahne, F., Hansen, K.D., Irizarry, R.A., Lawrence, M., Love, M.I., Macdonald, J., Obenchain, V., Oles, A.K., Pages, H., Reyes, A., Shannon, P., Smyth, G.K., Tenenbaum, D., Waldron, L., Morgan, M., 2015. Orchestrating high-throughput genomic analysis with Bioconductor. Nature Methods 12, 115–121. 10.1038/nmeth.3252

Jeffroy, F., Brulle, F., Paillard, C., 2013. Differential expression of genes involved in immunity and biomineralization during Brown Ring Disease development and shell repair in the Manila clam, Ruditapes philippinarum. Journal of Invertebrate Pathology 113, 129–136. 10.1016/j.jip.2013.03.001

Katsumiti, A., Gilliland, D., Arostegui, I., Cajaraville, M.P., 2014. Cytotoxicity and cellular mechanisms involved in the toxicity of CdS quantum dots in hemocytes and gill cells of the mussel Mytilus galloprovincialis. Aquatic Toxicology 153, 39–52. 10.1016/j.aquatox.2014.02.003

Li, X.S., Yang, H.L., Zhang, D.Y., Zhang, Y.M., Wood, A.J., 2012. Reference gene selection in the desert plant Eremosparton songoricum. International Journal of Molecular Science 13, 6944–6963. 10.3390/ijms13066944

Liu, N., Pan, L., Gong, X., Tao, Y., Hu, Y., Miao, J., 2014. Effects of benzo(a)pyrene on differentially expressed genes and haemocyte parameters of the clam Venerupis philippinarum. Ecotoxicology 23, 122–132. 10.1007/s10646-013-1157-7

Liu, X., Ji, C., Zhao, J., Wu, H., 2013. Differential metabolic responses of clam Ruditapes philippinarum to Vibrio anguillarum and Vibrio splendidus challenges. Fish & Shellfish Immunology 35, 2001–2007. 10.1016/j.fsi.2013.09.014

Mateo, D.R., Greenwood, S.J., Araya, M.T., Berthe, F.C., Johnson, G.R., Siah, A., 2010. Differential gene expression of gamma-actin, Toll-like receptor 2 (TLR-2) and interleukin-1 receptor-associated kinase 4 (IRAK-4) in Mya arenaria haemocytes induced by in vivo infections with two Vibrio splendidus strains. Developmental & Comparative Immunology 34, 710–714. 10.1016/j.dci.2010.02.006

Miao, J., Pan, L., Zhang, W., Liu, D., Cai, Y., Li, Z., 2014. Identification of differentially expressed genes in the digestive gland of manila clam Ruditapes philippinarum exposed to BDE-47. Comparative Biochemistry and Physiology C: Pharmacology Toxicology and Endocrinology 161, 15–20. 10.1016/j.cbpc.2013.12.004

Milan, M., Pauletto, M., Patarnello, T., Bargelloni, L., Marin, M.G., Matozzo, V., 2013. Gene transcription and biomarker responses in the clam Ruditapes philippinarum after exposure to ibuprofen. Aquatic Toxicology 126, 17–29. 10.1016/j.aquatox.2012.10.007

Moreira, R., Balseiro, P., Romero, A., Dios, S., Posada, D., Novoa, B., Figueras, A., 2012. Gene expression analysis of clams Ruditapes philippinarum and Ruditapes decussatus following bacterial infection yields molecular insights into pathogen resistance and immunity. Developmental & Comparative Immunology 36, 140–149. 10.1016/j.dci.2011.06.012

Moreira, R., Pereiro, P., Costa, M.M., Figueras, A., Novoa, B., 2015. Evaluation of reference genes of Mytilus galloprovincialis and Ruditapes philippinarum infected with three bacteria strains for gene expression analysis. Aquatic Living Resources 27, 147–152. 10.1051/alr/2014015

Morga, B., Arzul, I., Faury, N., Renault, T., 2010. Identification of genes from flat oyster Ostrea edulis as suitable housekeeping genes for quantitative real time PCR. Fish & Shellfish Immunology 29, 937–945. 10.1016/j.fsi.2010.07.028

Perkins, J.R., Dawes, J.M., Mcmahon, S.B., Bennett, D.L., Orengo, C., Kohl, M., 2012. ReadqPCR and NormqPCR: R packages for the reading, quality checking and normalisation of RT-qPCR quantification cycle (Cq) data. BMC Genomics 13, 296. 10.1186/1471-2164-13-296

Pfaffl, M.W., Tichopad, A., Prgomet, C., Neuvians, T.P., 2004. Determination of stable housekeeping genes, differentially regulated target genes and sample integrity: BestKeeper--Excel-based tool using pair-wise correlations. Biotechnology Letters 26, 509–515. 10.1023/B:BILE.0000019559.84305.47

R Core Team, 2016. R: A Language and Environment for Statistical Computing. R Foundation for Statistical Computing, Vienna, Austria.

Snape, J.R., Maund, S.J., Pickford, D.B., Hutchinson, T.H., 2004. Ecotoxicogenomics: the challenge of integrating genomics into aquatic and terrestrial ecotoxicology. Aquatic Toxicology 67, 143–154. 10.1016/j.aquatox.2003.11.011

Snell, T.W., Brogdon, S.E., Morgan, M.B., 2003. Gene expression profiling in ecotoxicology. Ecotoxicology 12, 475–483. 10.1023/B:ECTX.0000003033.09923.a8

Thellin, O., Zorzi, W., Lakaye, B., De Borman, B., Coumans, B., Hennen, G., Grisar, T., Igout, A., Heinen, E., 1999. Housekeeping genes as internal standards: use and limits. Journal of Biotechnology 75, 291–295. 10.1016/s0168-1656(99)00163-7

Thomas Reuters, 2016. 2015 Journal Citation Reports^®^.

Tricarico, C., Pinzani, P., Bianchi, S., Paglierani, M., Distante, V., Pazzagli, M., Bustin, S.A., Orlando, C., 2002. Quantitative real-time reverse transcription polymerase chain reaction: normalization to rRNA or single housekeeping genes is inappropriate for human tissue biopsies. Analytical Biochemistry 309, 293–300. 10.1016/s0003-2697(02)00311-1

Untergasser, A., Cutcutache, I., Koressaar, T., Ye, J., Faircloth, B.C., Remm, M., Rozen, S.G., 2012. Primer3--new capabilities and interfaces. Nucleic Acids Research 40, e115. 10.1093/nar/gks596

Vandesompele, J., De Preter, K., Pattyn, F., Poppe, B., Van Roy, N., De Paepe, A., Speleman, F., 2002. Accurate normalization of real-time quantitative RT-PCR data by geometric averaging of multiple internal control genes. Genome Biology 3, RESEARCH0034. 10.1186/gb-2002-3-7-research0034

Veldhoen, N., Lowe, C.J., Davis, C., Mazumder, A., Helbing, C.C., 2009. Gene expression profiling in the deep water horse mussel Modiolus modiolus (L.) located near a marine municipal wastewater outfall. Aquatic Toxicology 93, 116–124. 10.1016/j.aquatox.2009.04.002

Volland, M., Hampel, M., Martos-Sitcha, J.A., Trombini, C., Martinez-Rodriguez, G., Blasco, J., 2015. Citrate gold nanoparticle exposure in the marine bivalve Ruditapes philippinarum: uptake, elimination and oxidative stress response. Environmental Science and Pollution Research 22, 17414–17424. 10.1007/s11356-015-4718-x

Wan, Q., Whang, I., Choi, C.Y., Lee, J.S., Lee, J., 2011. Validation of housekeeping genes as internal controls for studying biomarkers of endocrine-disrupting chemicals in disk abalone by real-time PCR. Comparative Biochemistry and Physiology C: Pharmacology Toxicology and Endocrinology 153, 259–268. 10.1016/j.cbpc.2010.11.009

Wickham, H., 2009. ggplot2: Elegant Graphics for Data Analysis. Springer-Verlag New York.

Winnebeck, E.C., Millar, C.D., Warman, G.R., 2010. Why does insect RNA look degraded? Journal of Insect Science (Ludhiana) 10, 159. 10.1673/031.010.14119

Wong, M.L., Medrano, J.F., 2005. Real-time PCR for mRNA quantitation. BioTechniques 39, 75–85. 10.2144/05391RV01

Xie, F., Xiao, P., Chen, D., Xu, L., Zhang, B., 2012. miRDeepFinder: a miRNA analysis tool for deep sequencing of plant small RNAs. Plant Molecular Biology 80, 75–84. 10.1007/s11103-012-9885-2

Xu, C., Pan, L., Liu, N., Wang, L., Miao, J., 2010. Cloning, characterization and tissue distribution of a pi-class glutathione S-transferase from clam (Venerupis philippinarum): Response to benzo[alpha]pyrene exposure. Comparative Biochemistry and Physiology C: Pharmacology Toxicology and Endocrinology 152, 160–166. 10.1016/j.cbpc.2010.03.011

